# Metabarcode and transcriptome datasets of *Pinus sylvestris* to assess fungal phyllosphere and disease dynamics

**DOI:** 10.64898/2026.05.14.725107

**Authors:** Beth Moore, Annika Perry, Sundeep Kaur, Bridget Crampton, Amir Gurung, Joan K Beaton, Joan E Cottrell, Jenni A Stockan, V Anne Smith, Jenny Morris, Pete E Hedley, Krisztian Nemeth, Hattie Barber, Stephen Cavers, Susan Jones

**Affiliations:** The James Hutton Institute, Dundee, UK; UK Centre for Ecology & Hydrology, Penicuik, UK; Forest Research, Alice Holt Lodge, Surrey, UK; School of Biology, University of St Andrews, UK

**Author notes:** Corresponding author: Beth Moore.

## Abstract

Understanding how host–microbiome interactions influence tree disease is critical for understanding forest resilience. Here, we present foliar microbiome ITS2 metabarcoding transcriptomic datasets from *Pinus sylvestris* to investigate susceptibility to Dothistroma needle blight (DNB), a globally important foliar disease caused by *Dothistroma septosporum*. We hypothesised that host genotype shapes foliar microbial communities and their interactions, thereby influencing disease outcomes. Samples were collected from a progeny–provenance field trial in the south of Scotland representing a broad spectrum of disease susceptibilities. The dataset comprises ITS2 metabarcoding samples from 200 genotypes across three timepoints and RNAseq samples from 48 genotypes across two timepoints. Sampling captured key stages of pathogen exposure and disease progression. Both standardised and bespoke protocols were used for nucleotide extraction, sequencing, and quality control, including multiple negative and positive controls. These datasets, available in the European Nucleotide Archive (project accession PRJEB88228), enable analysis of temporal dynamics in foliar fungal communities, host–microbiome transcriptional responses, and genotype-dependent variation in disease susceptibility.

## Background & Summary

A key challenge in tree disease research is to understand how interactions between the host and its microbiome affect disease incidence and severity. *Dothistroma septosporum* is an ascomycete fungal pathogen that causes Dothistroma needle blight (DNB) in more than one hundred Pinaceae taxa worldwide^1^. DNB is a foliar disease, causing red-brown lesions on needles and premature needle loss. These symptoms result in reduced timber yields and, in severe cases, tree death. Scots pine (*Pinus sylvestris*), the most widely distributed pine species, is of great importance both ecologically and economically. *P*.*sylvestris* is susceptible to infection by *D*.*septosporum* and susceptibility is linked to genetic variability^2^. To test our hypothesis that host genotype drives foliar microbiome composition and interactions to alter host susceptibility to DNB we generated two datasets from a *P. sylvestris* progeny-provenance field trial in the south of Scotland, in which annual disease surveys had been conducted and trees genotyped ^3,4,5^. The two datasets generated were: (1) Fungal ITS2 metabarcode sequences of the foliar microbiota across three time points (July, August and September; during the DNB growing season) and (2) RNA sequences (RNAseq) of the foliar microbiota and the pine tree host across two time points (August and September) (Figure 1). The dataset comprised 585 samples from 200 genotypes across 3 time points for the ITS2 metabarcoding and 96 samples from 48 genotypes across 2 timepoints for the RNAseq. The tree genotypes sampled were designed to represent the full range of disease susceptibilities^4^, allowing the data to be used to analyse differences in foliar microbiomes across trees with different responses to DNB infection.

**Figure 1:**
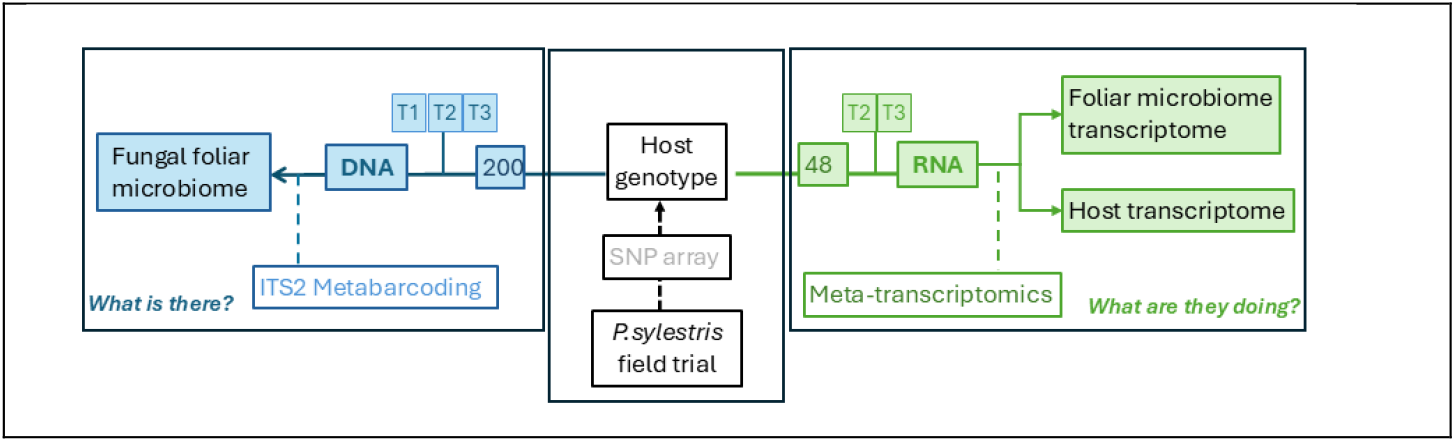
Overview diagram of the experiment and the datasets generated. Tree genotyping was conducted previously ^5^ and the diagram indicates the number of genotypes sampled for each part of the current foliar microbiome experiment. T1,T2 and T3 indicate the three timepoints for sample collection (July, August and September respectively).

## Methods

### Trial site

The host *P. Sylvestris* trees were located within a progeny-provenance trial in the south of Scotland, UK (FS, latitude 55.603625, longitude −2.893025). This site is part of a long-term multisite common garden trial and details of the trial design, and the phenotypic characterisation of the trees, have been published previously^3^. Briefly, the trial comprises a replicated block design with one individual from each of eight families per 21 populations (168 trees) represented in each of the four blocks^3^. A subset of these trees was selected for the current foliar microbiome study. In total, ten populations and five families within each population were selected (a total of 50 families) with four half-siblings included per family (total 200 trees) (one per sampling block). Annual disease surveys, conducted since 2015 for all trees in the trial, were used to characterise susceptibility to DNB across each population^4^ and ensured the selected trees represented the full range of disease susceptibilities.

### Collection of needle samples

Uninfected needles (only those that were visually inspected as green and healthy with no apparent lesions or discolouration) were collected at three timepoints during summer 2023.

- Timepoint 1 (TP1) (4-5 July) was shortly after budburst when needles were relatively newly emerged and exposure to the pathogen was expected to be relatively low
- Timepoint 2 (TP2) (15-16 August) was six weeks later, once the DNB infection cycle was expected to have progressed and host exposure to the pathogen increased
- Timepoint 3 (TP3) (27-28 September) was another six weeks later, when the DNB infection cycle was expected to have peaked for that year.

To reduce the influence of sampling position within the tree, two needle pairs from each of the cardinal directions (or as close as possible) and at a height of 1-2m were collected. Branches sampled in TP1 were labelled to ensure sampling was from the same location at subsequent timepoints. Sterile gloves and facemasks were used to limit contamination of samples. For the metabarcoding work, the needles were collected at the trial site into sterile plastic bags on ice and then transferred to a -20°C freezer for storage prior to DNA extraction. For metabarcoding, needles were collected at all three time points (TP1, TP2 and TP3). For transcriptomics, the needles were collected at the trial site into sterile plastic bags and immediately snap-frozen using dry ice. They were transferred from the trial site to the sequencing laboratory on dry ice and then stored in a -80°C freezer prior to RNA extraction. For transcriptomics, needles were only collected at two timepoints (TP2 and TP3) when the DNB transcriptome should be detectable.

### DNA extraction and metabarcode sequencing

Needles were freeze-dried, homogenized and total cellular DNA extracted using the Qiagen DNeasy Plant 96 Kit under standard protocol^6^, except for the addition of 10 µl proteinase K per sample during the incubation stage^6^. Amplicon library preparation was conducted using standard Illumina 16S metagenomic library preparation protocol^7^, with the exception that the number of PCR cycles were increased (PCR conditions: 95 °C for 3 min; 40 cycles of 95 °C for 30s, 55 °C for 30s, 72 °C for 30s; final extension at 72 °C for 5 min). Two primers were used to amplify the fungal ITS2 region

- fITS7 [TCGTCGGCAGCGTCAGATGTGTATAAGAGACAG]GTGARTCATCGAATCTTTG
- ITS4r [GTCTCGTGGGCTCGGAGATGTGTATAAGAGACAG]TCCTCCGCTTATTGATATGC

Illumina IDT unique dual indexes^8^ were used to barcode the samples (PCR conditions: 95 °C for 3 min; 8 cycles of 95 °C for 30s, 55 °C for 30s, 72 °C for 30s; final extension at 72 °C for 5 min) and the resulting libraries were quality checked on a Nanodrop (LabTech). In addition, a random subset of samples was checked using a Bioanalyzer 2100 (Agilent). Samples were normalised to a concentration of 8 nM before being pooled for sequencing.

Library pool quality control was performed using Qubit 2 fluorometer, qPCR (NEB kit E7630L) on StepOnePlus (Thermo Fisher), and Bioanalyzer 2100 to check library concentration and fragment size respectively. Pooled library (12 µl) and 8 µl of PhiX were combined and loaded at 1000 pM for sequencing. A total of six pools were sequenced on a NextSeq 2000 at the James Hutton Institute using a 2 × 300 bp paired end strategy.

### Metabarcoding controls

Negative controls were included at multiple stages of library preparation. DNA extraction blanks (DNA-blank) comprised lysis buffer without the addition of any DNA material and were carried through to the rest of the pipeline. Amplicon and index blanks comprised elution buffer (instead of eluted DNA) added from the start of library preparation (AMP-blank) and before the addition of Illumina indexes (IDX-blank). Positive microbial community controls were also used. A ZymoBIOMICS® whole cell Microbial Community Standard (catalogue no. D6300 ^9^) was included from the DNA extraction stage of each plate (SYN-extr)and three ZymoBIOMICS® Microbial Community DNA Standards (catalogue no. D6305)^10^ (SYN-rep) were included at the start of library prep. Controls were randomly allocated across plates with a single DNA-blank and SYN-extr control included at the DNA extraction stage. An additional blank (AMP-blank) was added to each plate at the start of library prep, along with four replicates of the Zymobiomics DNA standard (SYN-Rep). A final blank (IDX-blank) was added to each plate prior to the index PCR step. In addition, tree genotype 7142 at timepoint 1 was selected as a technical replicate and the single DNA extraction for this sample included three times on each plate. The numbers of each control type at each time point are summarized in Table 1.

**Table 1:** Numbers of different control types included for the ITS2 metabarcode sequencing at the three time points (T1:July, T2:August, T3:September)

| Control_type | Description | T1 | T2 | T3 | Total |
| --- | --- | --- | --- | --- | --- |
| Amp-Blank | Amplicon blank | 1 | 2 | 2 | 5 |
| DNA-Blank | DNA extraction blanks | 1 | 2 | 3 | 6 |
| IDX-Blank | Index blank | 1 | 1 | 2 | 4 |
| Pine-Rep | Technical rep: genotype 7142 T1 | 8 | 9 | 9 | 26 |
| Syn-Extr | Synthetic community extraction | 2 | 3 | 3 | 8 |
| Syn-Rep | Synthetic community DNA standard | 12 | 12 | 12 | 36 |

### Bioinformatics processing of metabarcode sequences

Reads were processed with fastp^11^ to remove short reads (<90 bp) and polyG tails. ITSxpress^12^ was used to remove flanking regions surrounding the target ITS2 sequence. Qiime2^13^ was then used for downstream analysis. Cutadapt^14^ was used within Qiime2 to detect and remove adapters. DADA2^15^ was used for denoising; removing reads with high error rates (`max_ee_f: 2, max_ee_r:2 per read`); merging forward and reverse reads (`min_overlap:12, max_merge_mismatch: 0`) and removal of chimeric sequences under default settings (method: concensus, `min_fold_parent_over_abundance:1`). Read quality was visually assessed with qiime2 viewer prior to DADA2 but no additional read trimming was conducted (`--p-trunc-len-f 0, --p-trunc-len-r 0`). A single operational taxonomic unit (OTU) table was created per timepoint using the UNITE database (version 9.0)^16^ with the qiime2 feature-classifier running classify-sklearn^13,17^.

The OTU tables for each timepoint were then filtered to remove OTUs derived from controls. OTUs identified in any negative controls (DNA-blanks, AMP-blanks and IDX-blanks) were removed. The two fungal taxa in the positive control communities were *Saccharomyces cerevisiae* and *Cryptococcus neoformans*. Any OTU within the Saccharomyces or Cryptococcus genus were also removed to give the primary filtered OTU tables (see Data records).

### RNA extraction and sequencing

We used a modified CTAB method based on Chang^18^ to extract total RNA from pine needles collected at timepoints TP2 and TP3. In summary, this involved grinding needles under liquid nitrogen, and CTAB extraction buffer at 65°C with β-mercaptoethanol added to lyse cells, followed by phase separation using chloroform:IAA. RNA was then selectively precipitated from the aqueous phase using LiCL and incubated overnight at 4°C. RNA pellets were then collected, washed with 70% ethanol, and dissolved in SSTE, followed by another chloroform:IAA extraction and ethanol precipitation. Finally, pellets are washed, dried and resuspended in water. The full protocol is provided in Supplementary information. The RNA extracted was then quality checked for concentration and purity with a Qubit (Thermo Fisher) and Nanodrop (LabTech) respectively, and integrity using a Bioanalyzer 2100 (Agilent). The RNA was then processed by Novogene (https://www.novogene.com/) using their standard stranded RNAseq pipeline, generating paired-end 150 bp data as fastq data files.

### Bioinformatics processing of RNAseq

Reads were quality filtered using fastp (v0.23.4)^11^ to remove poly tails and low quality reads `(--trim_poly_x, --length_required 50, --qualified_quality_phred 20`). Ribosomal RNA was removed using sortmeRNA (v4.3.6)^19^ with the v4.3 sensitive database. Identification of host and non-host tRNA was performed using Hisat2 (v2.2.1)^20^. An index was built using the *P. tabuliformis* genome sequence^21^, which is the most closely related genome available for *P. sylvestris*. The quality filtered reads were mapped against this genome (--score-min “L,0,-0.2”) and mapped reads were defined as the host RNAseq dataset. A second enhanced index was built with *P. tabuliformis*, PhiX (NCBI RefSeq: GCF_000819615.1), *P. sylvestris* mitochondrial (GenBank: KY302806.1) and *P. sylvestris* chloroplast (GenBank: KR476379.1) genomes. The quality filtered reads were mapped to this enhanced genome index (--score-min “L,0, -0.2”, --un-conc-gz) and reads that failed to map were defined as the non-host RNAseq dataset.

### Data Records

The raw metabarcode sequences and raw RNAseq reads were deposited into the European Nucleotide archive: https://www.ebi.ac.uk/ena/browser/view/PRJEB88228. Supplementary data table S1 and S2 provides an overview of the samples and read counts in the deposited datasets.

The primary filtered OTU tables (one per time point) calculated from the ITS2 metabarcoding sequences are available on GitHub (https://github.com/HuttonICS/PineBiomeDataPaper, https://doi.org/10.5281/zenodo.20179422).

### Data Overview

#### Metabarcode sequences

A summary of the ITS read counts for all 592 samples across the three timepoints are shown in supplementary table S1. This table shows counts per sample for raw, filtered, denoised, merged, non-chimeric and non-contaminant reads. There were 7 samples that were removed from subsequent the analysis. Four samples had no raw sequenced reads (AC1-7050-T1-ITS, GT5-7253-T1-ITS, AC2-7291-T3-ITS, AC2-7474-T3-ITS) and one sample had no sequenced reads that survived the quality filtering process (AC9-7142-T1-ITS). Two samples were outliers (GE1-7112-T1-ITS had 14 times the median number of raw reads and CR3-7338-T3-ITS had an outlying taxonomic profile relative to other tree samples likely indicating an issue in library preparation or sequencing of this sample. Plots of these outliers are available in the github repository (https://github.com/HuttonICS/PineBiomeDataPaper/Plots). The raw read count per sample (excluding removed samples) ranged from 53,115 to 840,255 with an average 331,566 reads across all 3 timepoints. The final read counts per samples after filtering and denoising ranged from 35 962 to 565,097 with an average of 194,241 reads across all time points.

### RNAseq

A summary table of the RNA read counts for all 96 samples across T2 and T3 timepoints are shown in supplementary table S2. This table shows counts per sample for raw, filtered, host and non-host reads. The filtered total read count per sample ranged from 43,630,667 to 73,430,067 with a median count of 50,010,306. The number of host reads ranged from 34,037,896 to 60,121,826 with an average of 40,105,360, with the percentage of host reads varying from 72.5%-83.3%.

### Technical Validation

A single genotype (genotype 7142) was used as a technical replicate for the ITS2 metabarcoding. Needles from this tree at timepoint 1 were used across all plates to allow analysis of any potential batch effects. Relative abundance of OTUs grouped by class in these technical replicates is presented in Figure 2.

**Figure 2.**
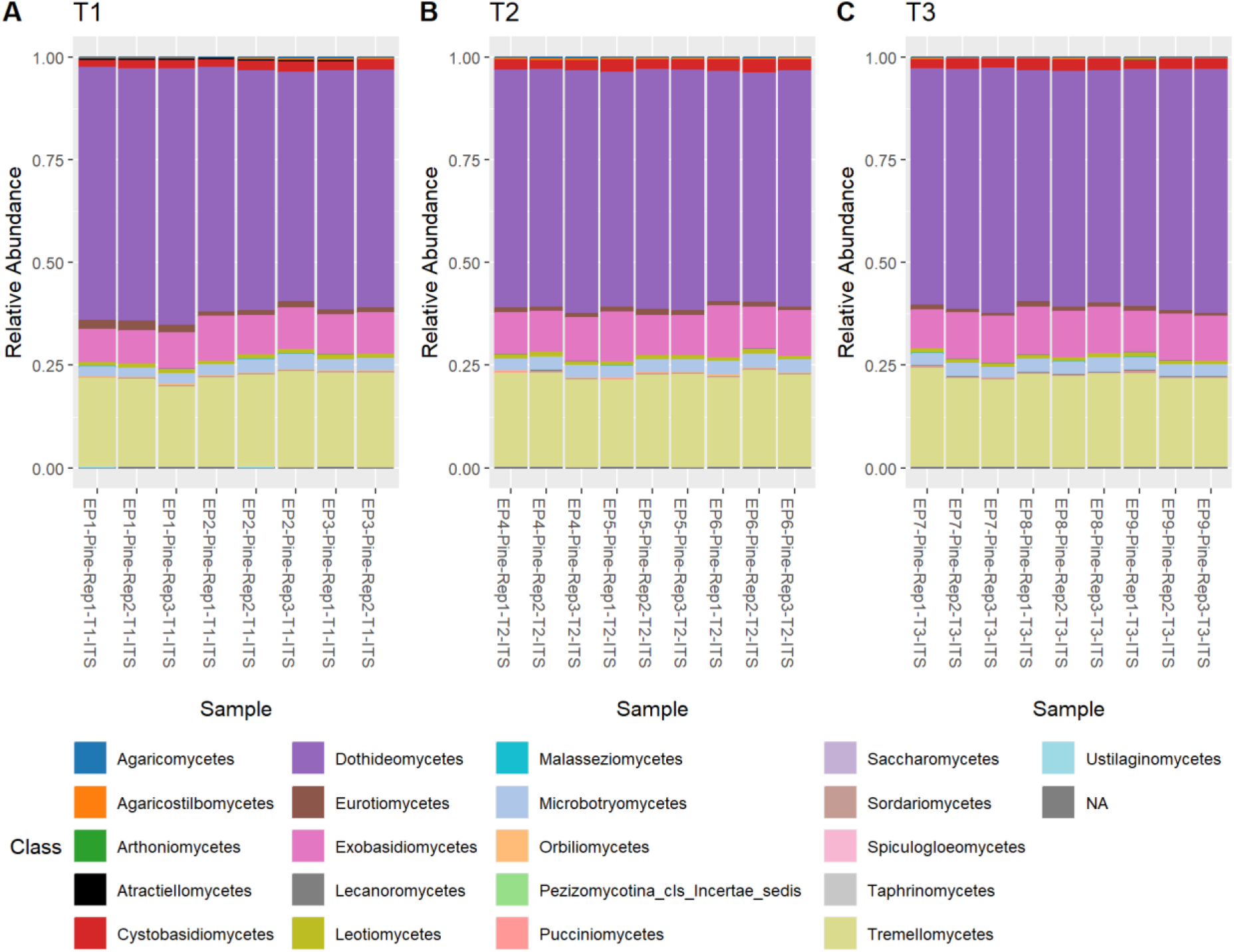
Relative abundance of taxa grouped by class for timepoints (A) T1, (B) T2, and (C) T3 for the technical replicates. Sample names indicate the extraction plate, with EP1 indicating a techincal replicate within plate 1.

Annual disease surveys allowed the ITS2 metabarcoding samples and the RNAseq samples to be split into three distinct groups, “low susceptibility”, “moderate susceptibility” and “high susceptibility” to disease^4^. Integrating the susceptibility group data with the fungal OTU tables allows the datasets to be used to assess genotype-dependent variation in disease susceptibility. Integrating the susceptibility group data with differential gene expression analysis allows the exploration of transcriptional variations linked to disease susceptibility in both host and foliar microbiome.

## Data Availability

Metabarcode sequences and RNA sequences are deposited in the ENA project PREJEB88228 (https://www.ebi.ac.uk/ena/browser/view/PRJEB88228). Supplementary tables providing an overview of the read counts throughout bioinformatic processing are available in supplementary information. Operational Taxonomic Unit tables are available from the github repository https://github.com/HuttonICS/PineBiomeDataPaper (https://doi.org/10.5281/zenodo.20179422).

## Code Availability

The code used to process the metabarcode sequence dataset and the RNAseq dataset is available on GitHub (https://github.com/HuttonICS/PineBiomeDataPaper).

## Acknowledgements

We thank Glenn Iason for the original pine trial establishment concept, planning and establishment and Carolyn Riddell for metabarcoding discussions.

## Author contributions

JB, JC and JS collected seed, cultivated seedlings and contributed to the multi-site common garden design and management. SJ, AP and SC designed the foliar microbiome experiments. SJ, BM, AP and SC wrote the manuscript. BM, AP, KN and HB conducted sample collection and preparation. SK, BC and AG created metabarcoding libraries. JM and BM conducted the RNA-sequence extractions and JM and PH conducted the RNA-sequencing. BM conducted all data analysis. Correspondence and requests for additional information should be addressed to BM and AP.

